## Supplementary Figures for "Transcription start sites experience a high influx of heritable variants fuelled by early development"

### Extended Data Figures

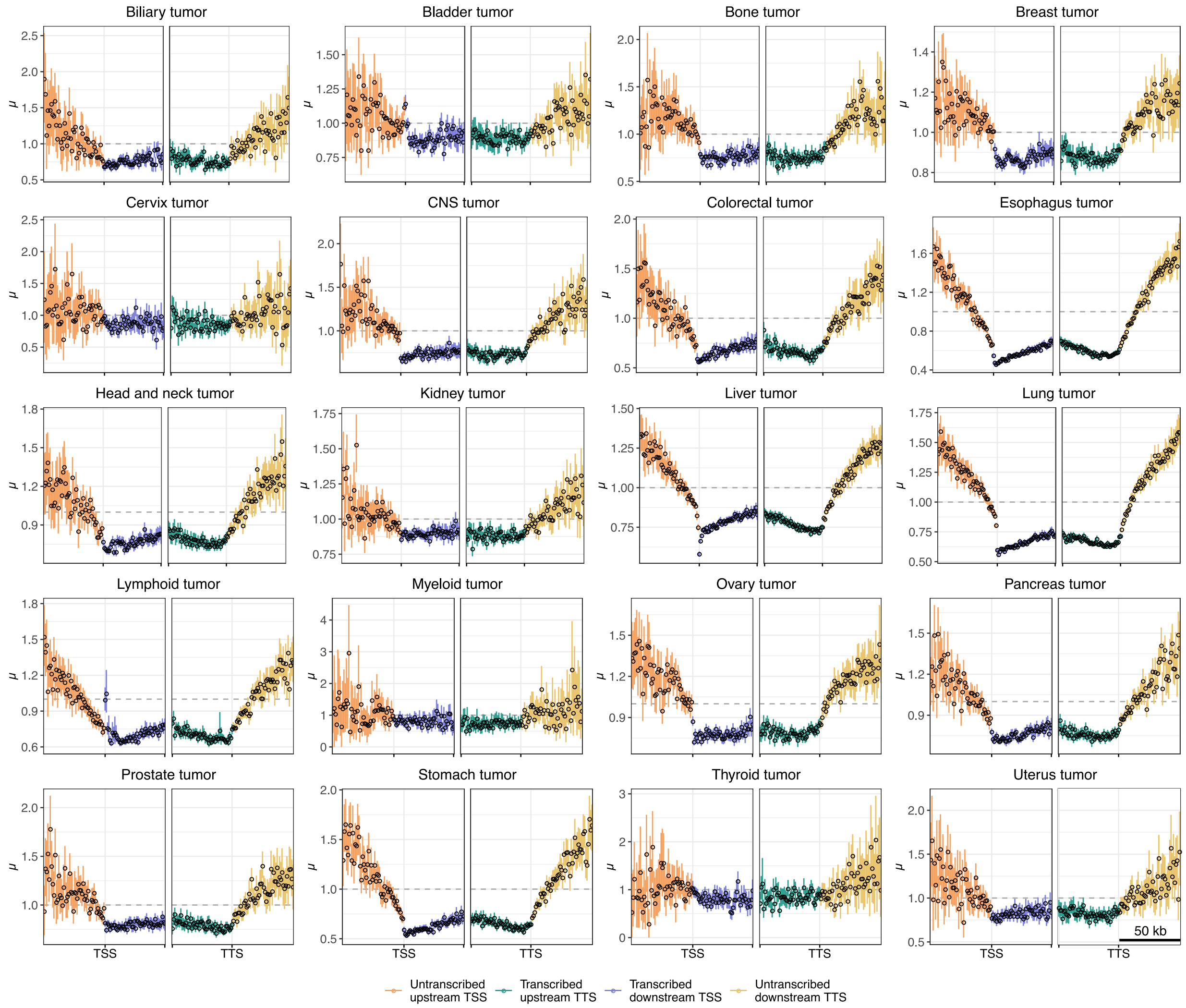


**Extended Data Figure 1:** Average number of non(CpG>TpG) mutations across 14763 protein-coding genes divided by the expectation based on the 5-mer sequence context, $\mu$, upstream and downstream of the TSS (orange and blue, respectively) and upstream and downstream of the TTS (green and yellow, respectively) in 1-kb windows. Error bars represent the 90% confidence intervals across 100 bootstrap replicates. Results are shown for all tumour types in the PCAWG dataset, indicated above each plot.

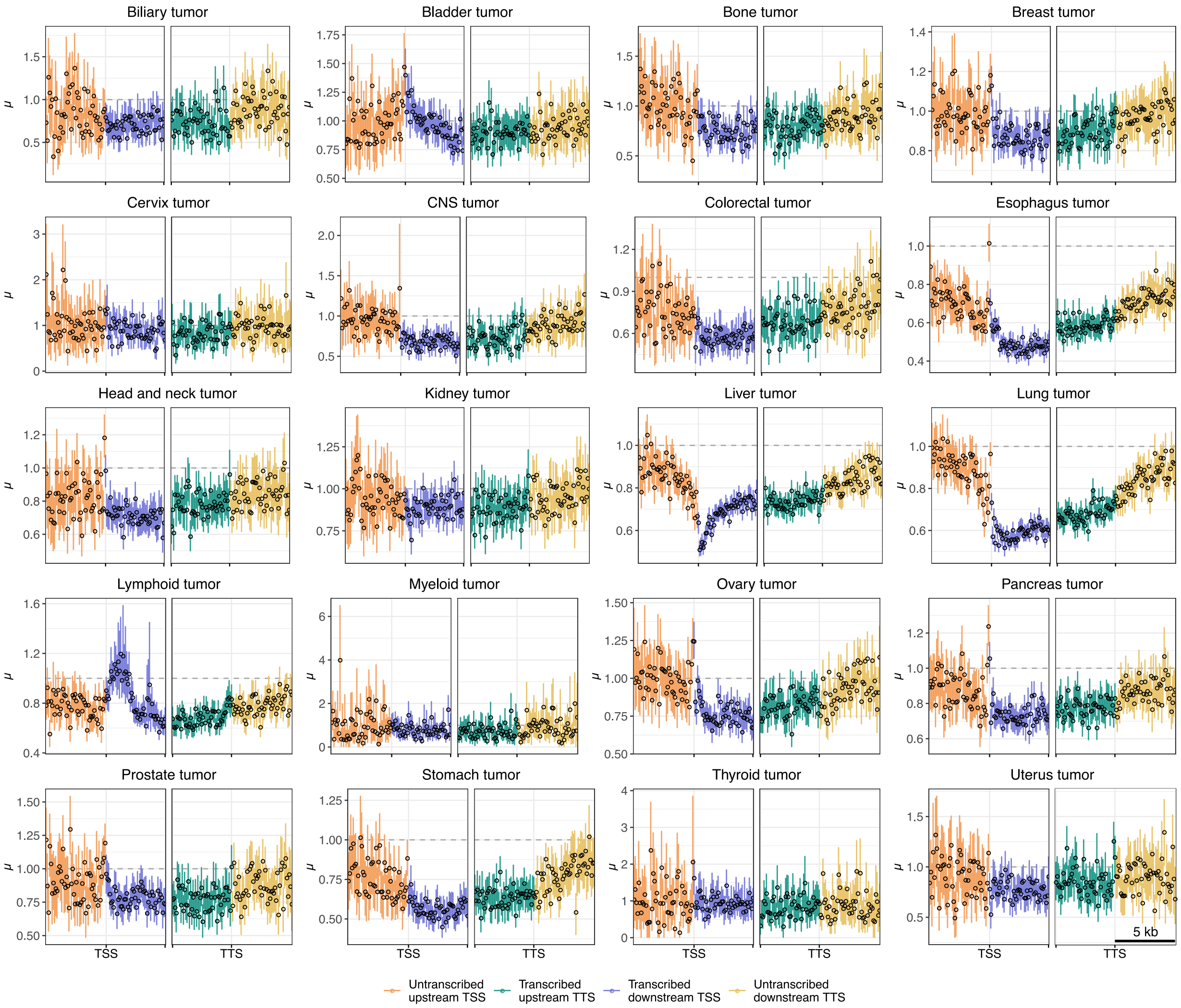


**Extended Data Figure 2:** Same as **Extended Data Figure 1**, but for 100-bp windows.

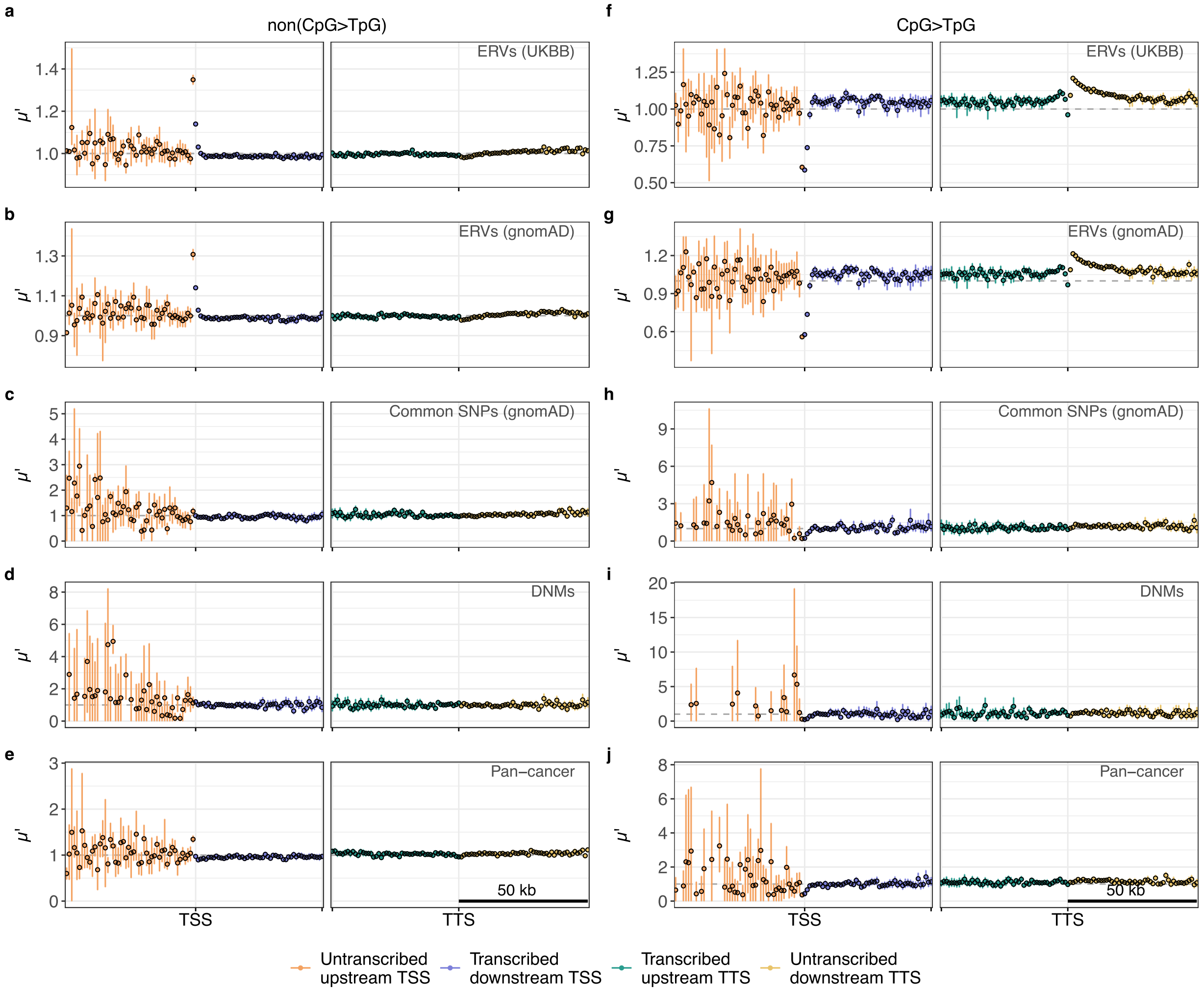


**Extended Data Figure 3:** Average number of (**a-e**) non(CpG>TpG) and (**f-j**) CpG>TpG mutations across 3991 divergent long non-coding RNA genes divided by the expectation based on the 5-mer sequence context and by the mean transcript-specific mutation density, $\mu'$, upstream and downstream of the TSS (orange and blue, respectively) and upstream and downstream of the TTS (green and yellow, respectively) in 1-kb windows. Error bars represent the 90% confidence intervals across 100 bootstrap replicates. Results are shown for (**a,f**) UKBB ERVs, (**b,g**) gnomAD ERVs, (**c,h**) common gnomAD SNPs (10%<AF<90%), (**d,i**) DNMs and (**e,j**) PCAWG pan-cancer mutations.

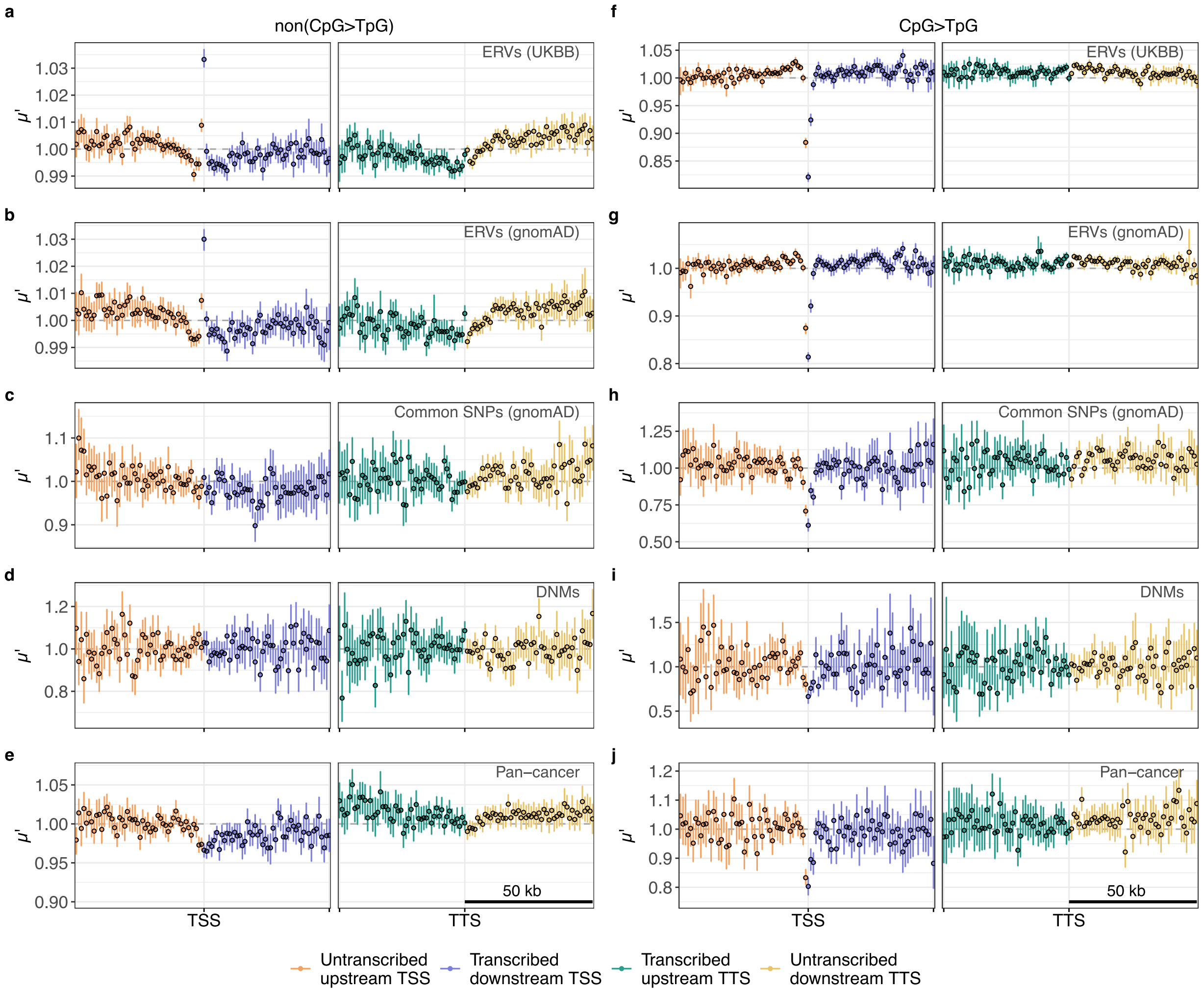


**Extended Data Figure 4:** $\mu'$ in 1-kb windows on and around 8454 intergenic long non-coding RNA genes. Panels same as in **Extended Data Figure 3**.

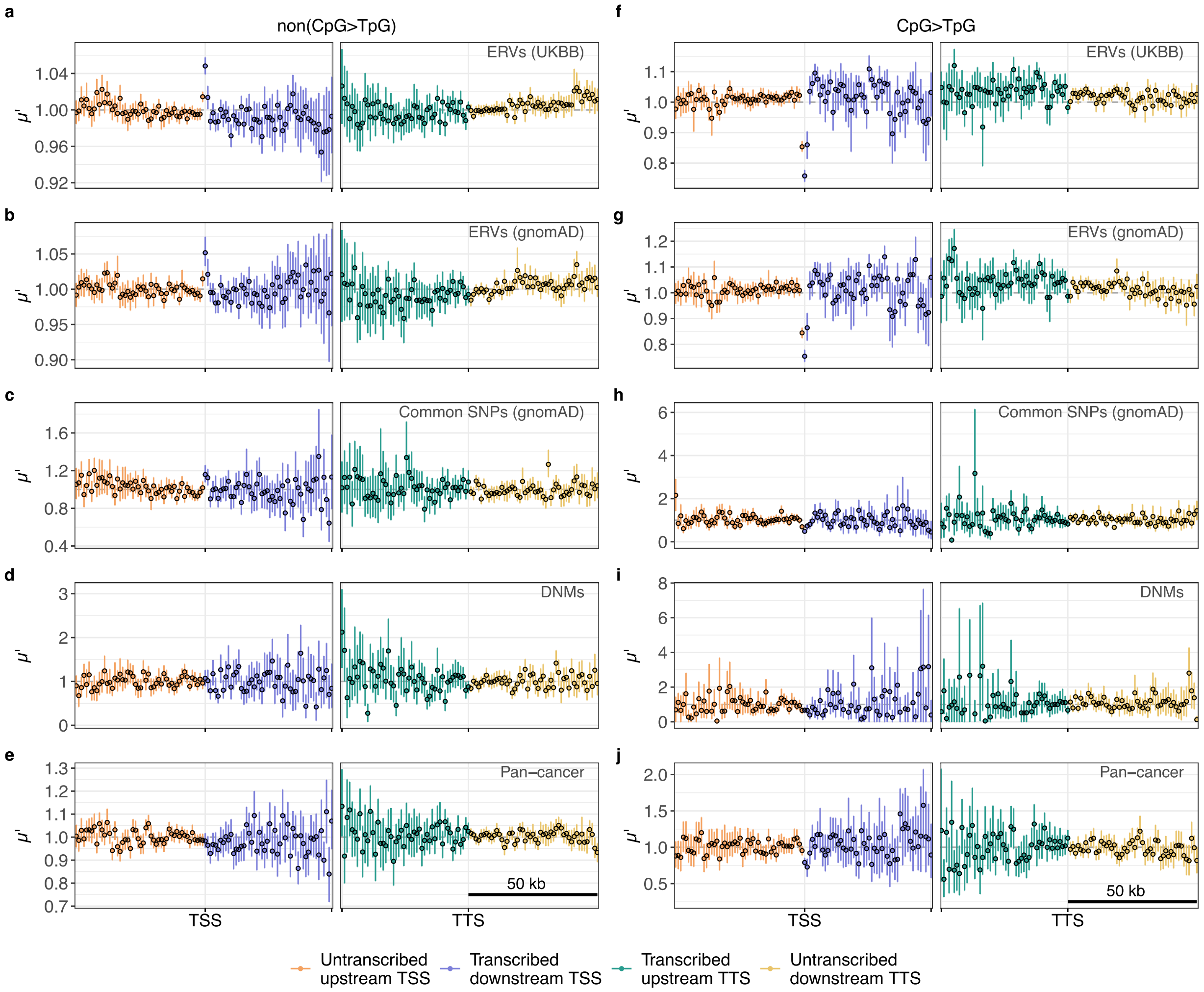


**Extended Data Figure 5:** $\mu'$ in 1-kb windows on and around 1660 pseudogenes. Panels same as in **Extended Data Figure 3**.

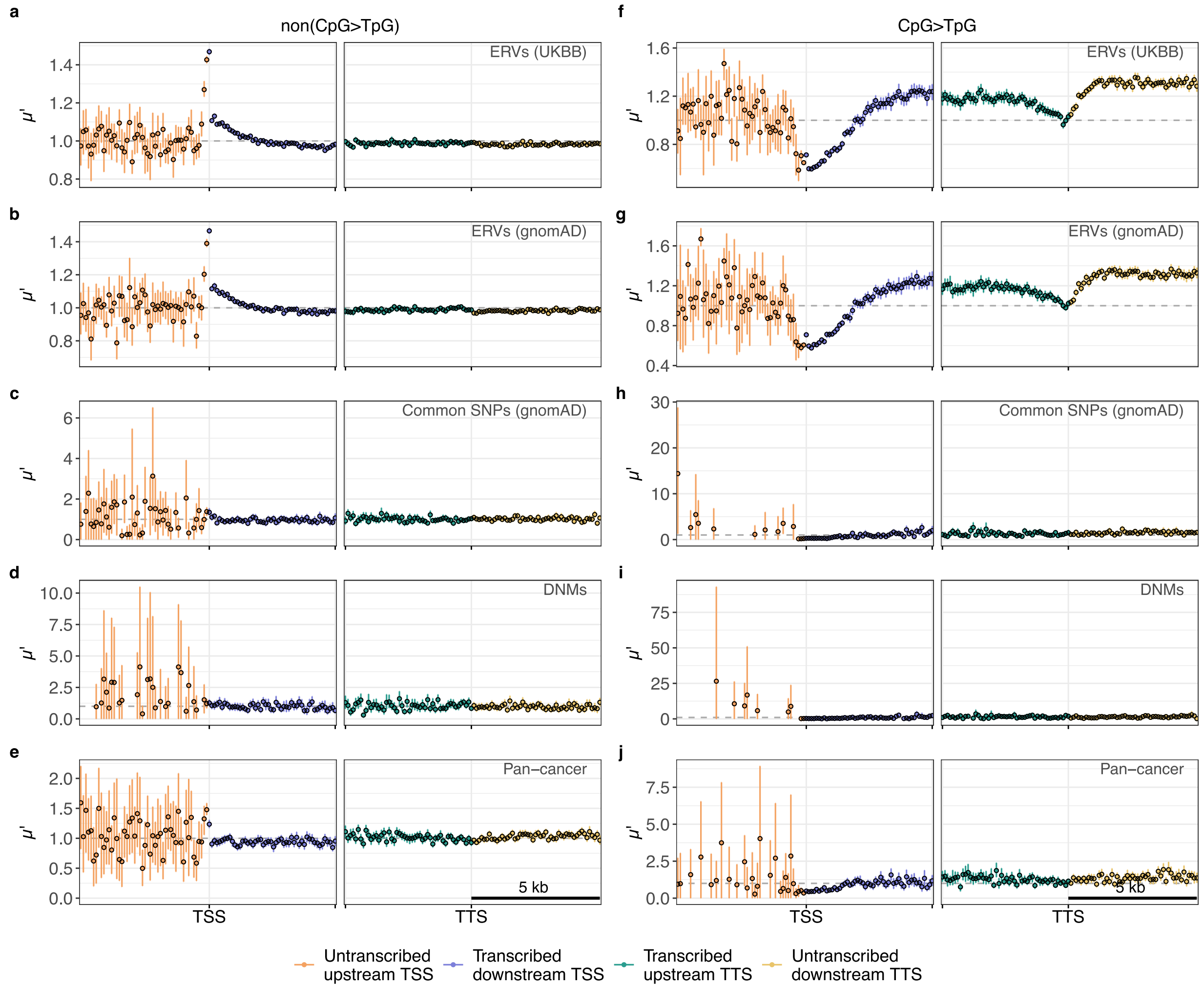


**Extended Data Figure 6:** $\mu'$ in 100-bp windows on and around 3991 divergent long non-coding RNA genes. Panels same as in **Extended Data Figure 3**.

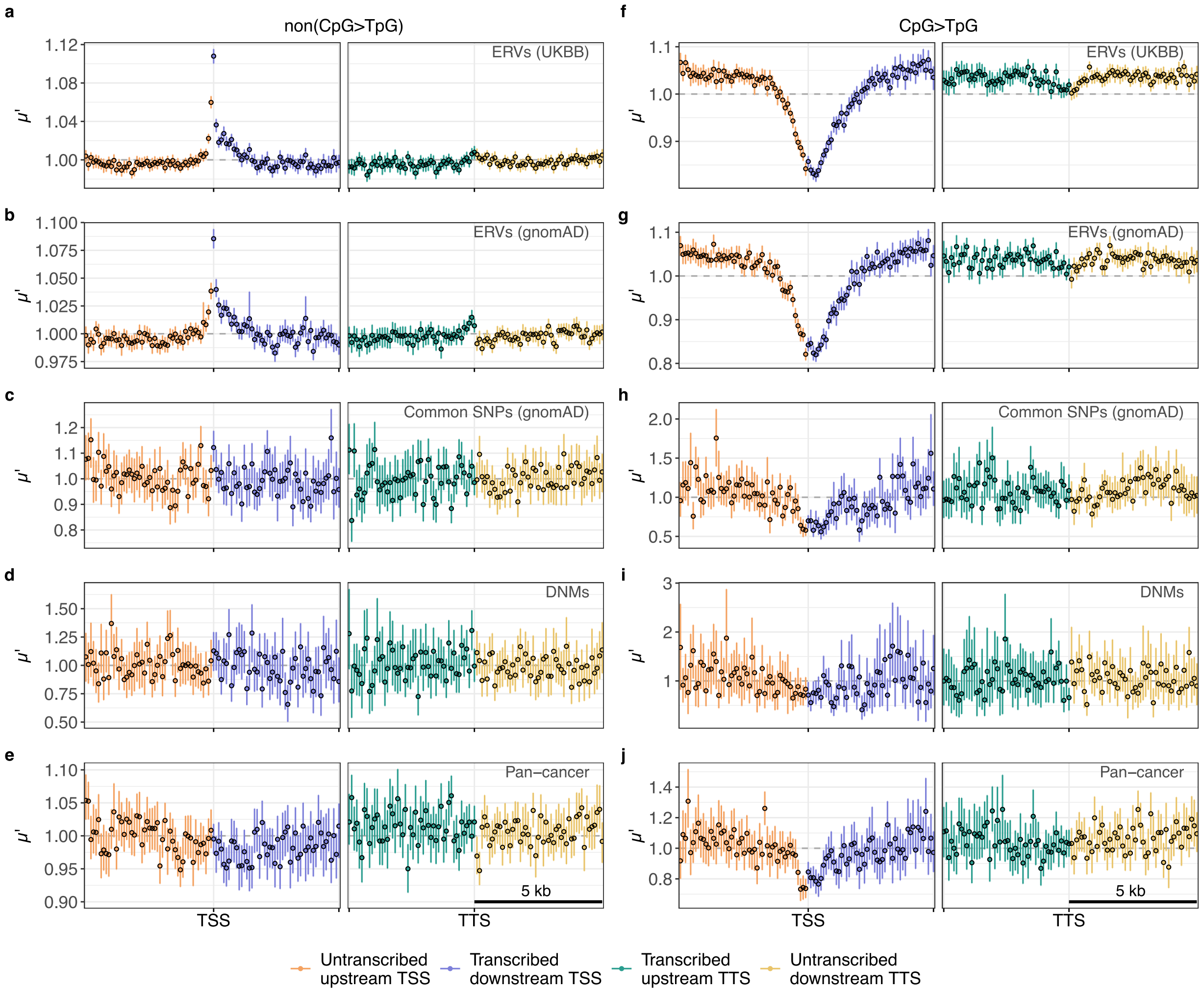


**Extended Data Figure 7:** $\mu'$ in 100-bp windows on and around 8454 intergenic long non-coding RNA genes. Panels same as in **Extended Data Figure 3**.

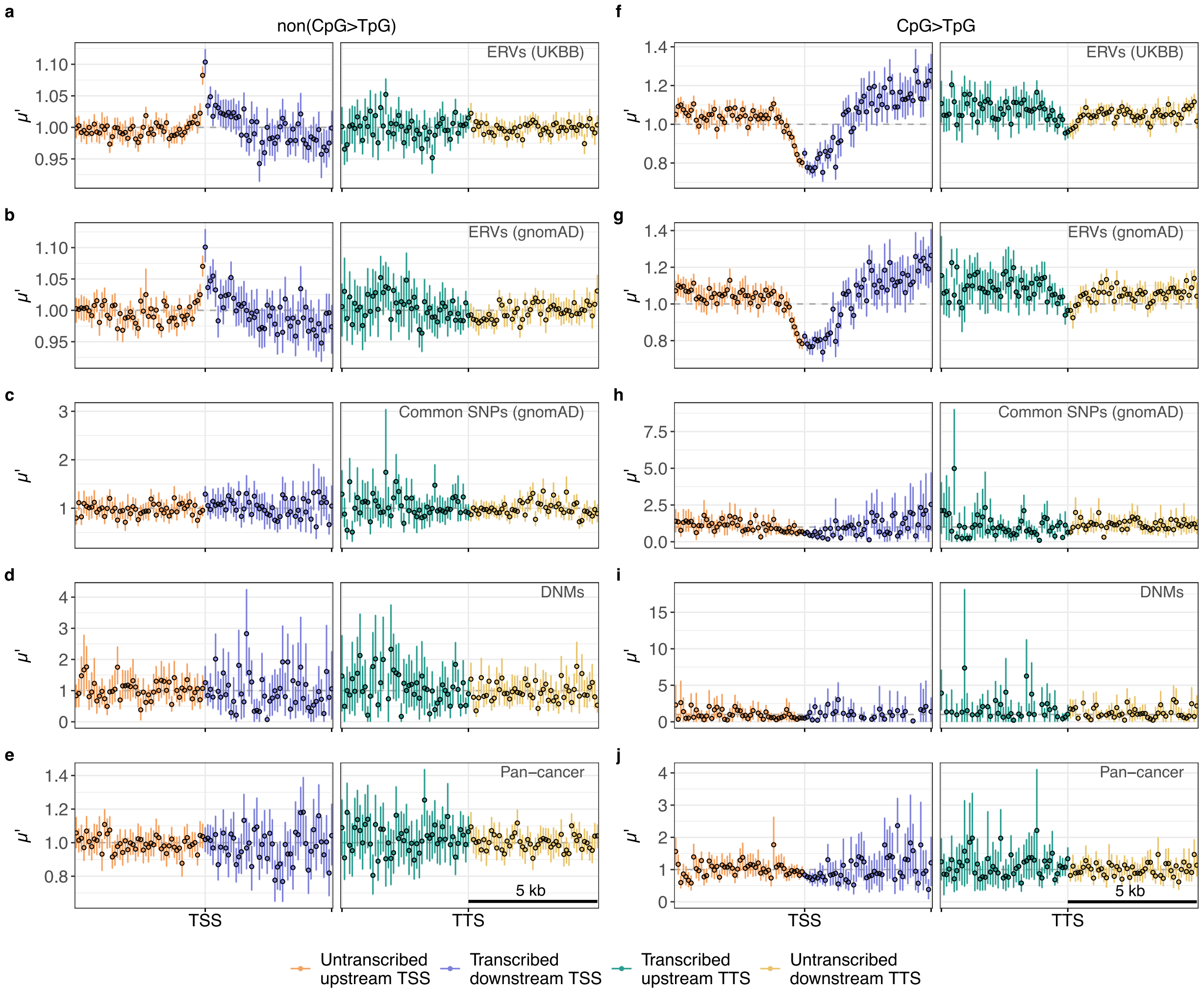


**Extended Data Figure 8:** $\mu'$ in 100-bp windows on and around 1660 pseudogenes. Panels same as in **Extended Data Figure 3**.

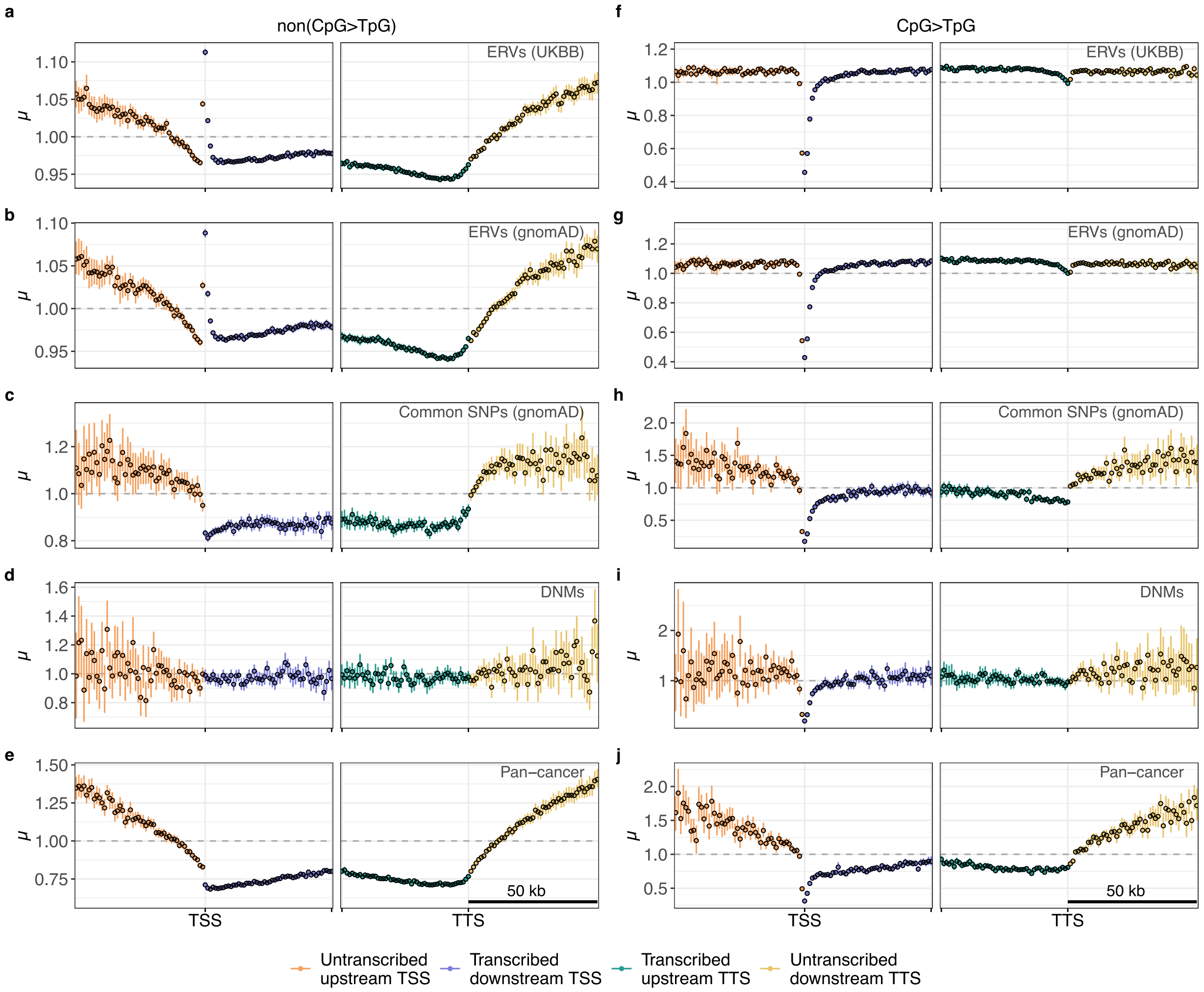


**Extended Data Figure 9:** $\mu$ in 1-kb windows on and around 14763 protein-coding genes. Panels same as in **Extended Data Figure 3**.

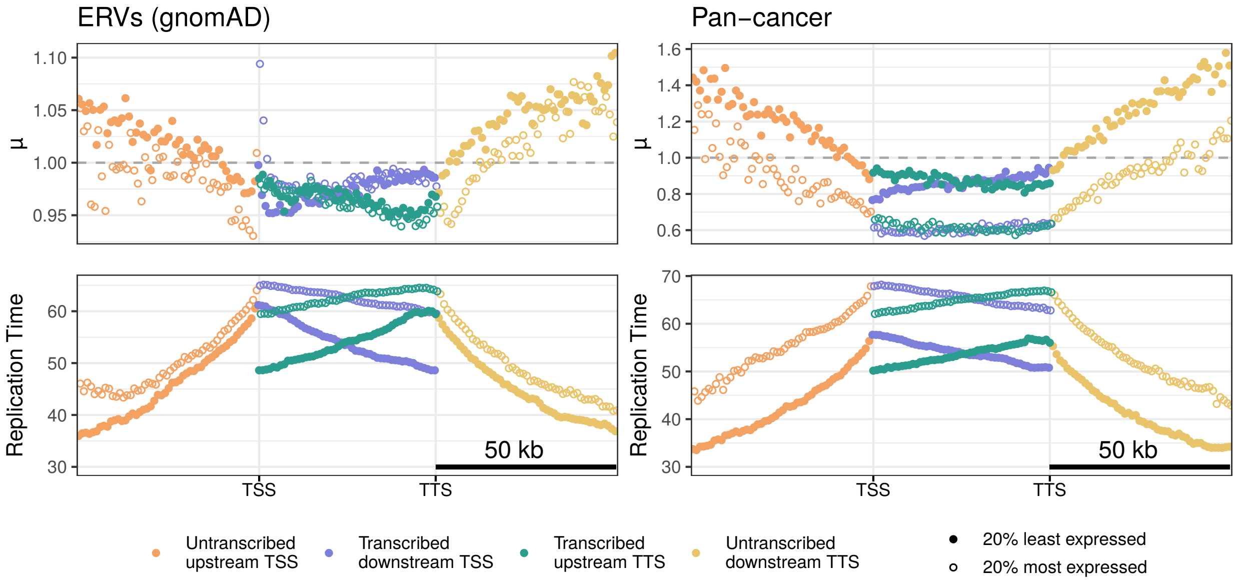


**Extended Data Figure 10: (a)** $\mu$ for the 20% most highly expressed and the 20% most lowly expressed protein-coding genes for ERVs (left) and pan-cancer (right). Expression levels from human testis were used for ERVs, while for pan-cancer the weighted mean over tissue-specific expression levels was calculated. **(b)** Replication time averaged over the two gene sets stratified by expression. The negative correlation between gene length and replication time (note that "early" corresponds to high and "late" to low values of replication time) together with the negative correlation between replication time and mutation rate cause the observed positive slope of $\mu$ on transcribed regions downstream of the TSS (blue) and a corresponding negative slope on transcribed regions upstream of the TTS (green). Division of each window-specific $\mu_{tb}$ by the mean of the transcript ($\mu_t$) produces $\mu'$.

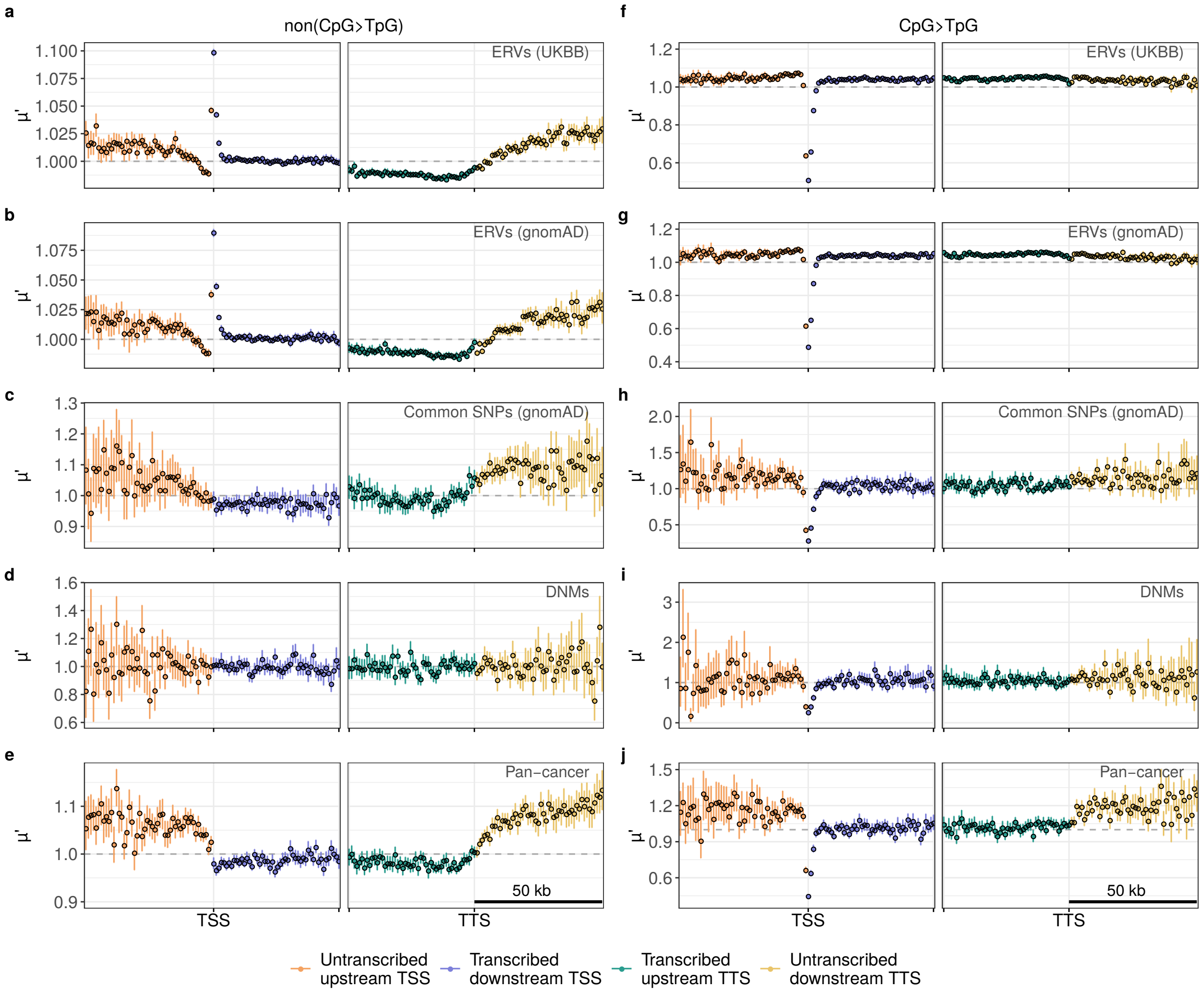


**Extended Data Figure 11:** $\mu'$ in 1-kb windows on and around 14763 protein-coding genes, with coding exons and conserved non-coding sequence elements excluded. Panels same as in **Extended Data Figure 3**.

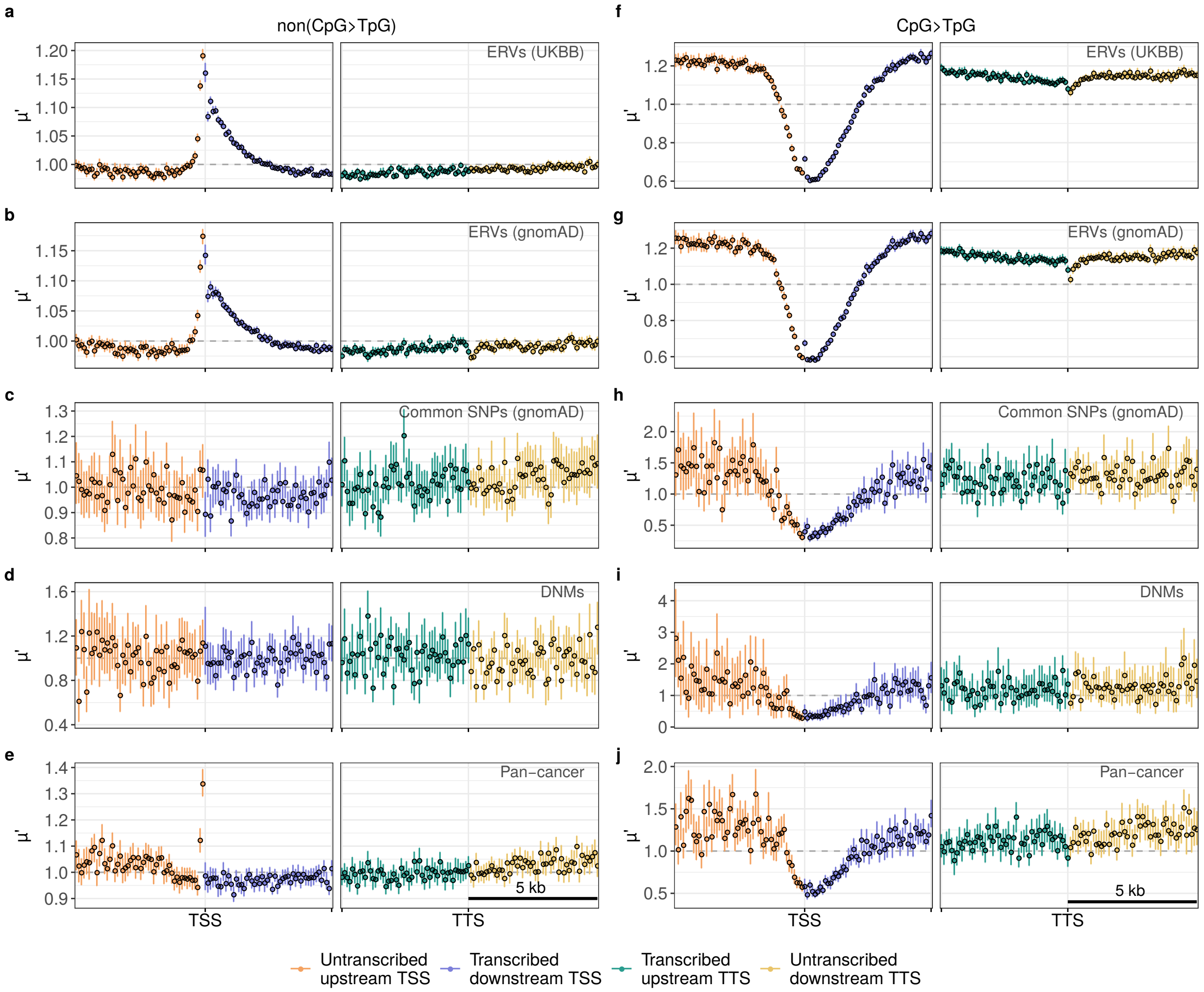


**Extended Data Figure 12:** $\mu'$ in 100-bp windows on and around 14763 protein-coding genes, with coding exons and conserved non-coding sequence elements excluded. Panels same as in **Extended Data Figure 3**.

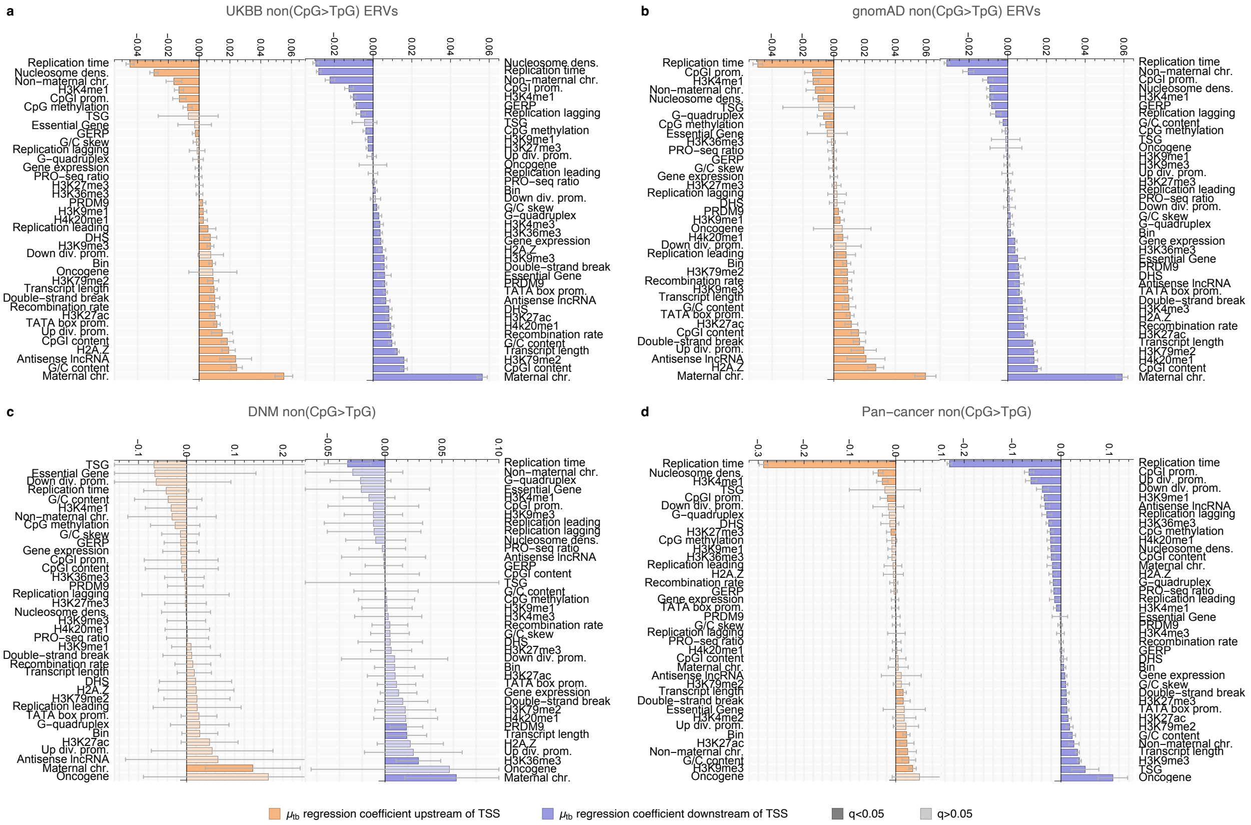


**Extended Data Figure 13:** Standardised regression coefficients from multiple negative binomial regression of $\mu_{tb}$ for non(CpG>TpG) mutations across all transcripts and 1-kb windows upstream of the TSS (orange) and downstream of the TSS (blue). Results are shown for **(a)** UKBB ERVs, **(b)** gnomAD ERVs, **(c)** DNMs and **(c)** PCAWG pan-cancer mutations. Note that "early" corresponds to high and "late" to low values of replication time. Error bars show multiple-testing-adjusted 95% confidence intervals. Dark shading indicates a multiple-testing-adjusted p-value (q-value) of less than 0.05.

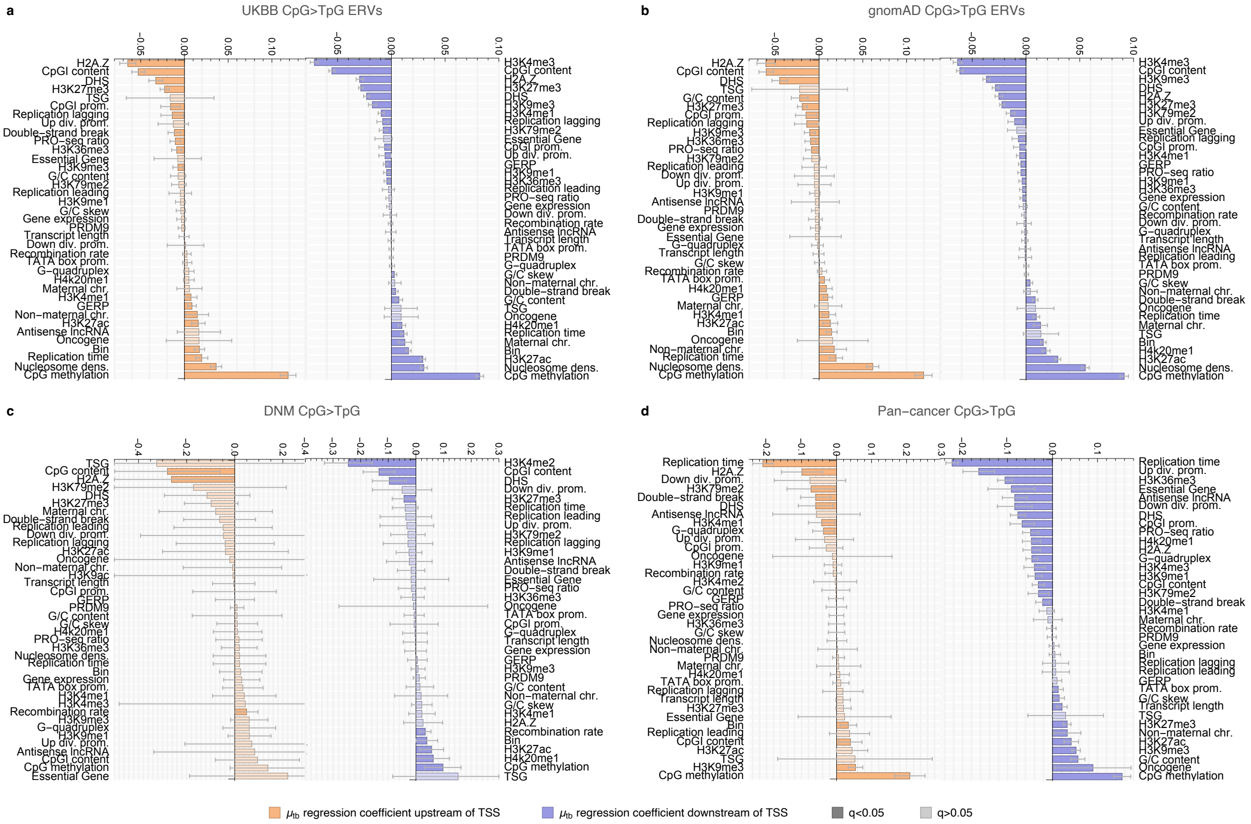


**Extended Data Figure 14:** Standardised regression coefficients from multiple Poisson regression of $\mu_{tb}$ for CpG>TpG mutations across all transcripts and 1-kb windows upstream of the TSS (orange) and downstream of the TSS (blue). Same panels as in **Extended Data Figure 13**.

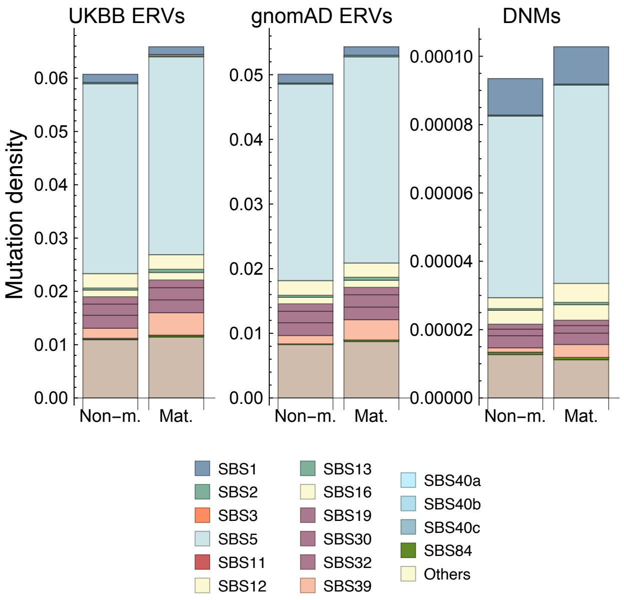


**Extended Data Figure 15:** Mutation density stratified by mutational signature from all considered genomic regions around genes on non-maternal chromosomes (1, 13, 18, 20) and maternal chromosomes (8, 9, 15, 16) for UKBB ERVs, gnomAD ERVs and DNMs (left to right), showing that SBS39 is increased on maternal chromosomes.

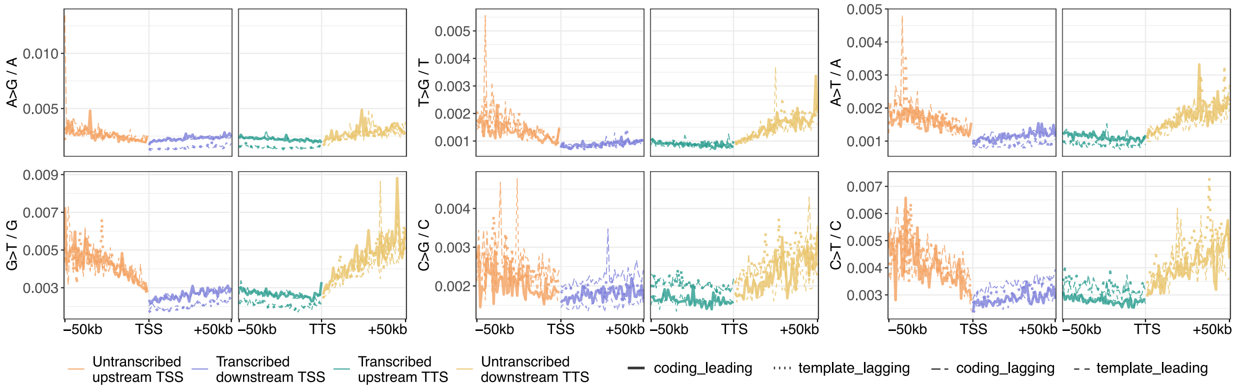


**Extended Data Figure 16:** non(CpG>TpG) pan-cancer mononucleotide mutation density, stratified by transcription strand (coding or template) and replication strand (leading or lagging) in 1-kb windows around the TSS.

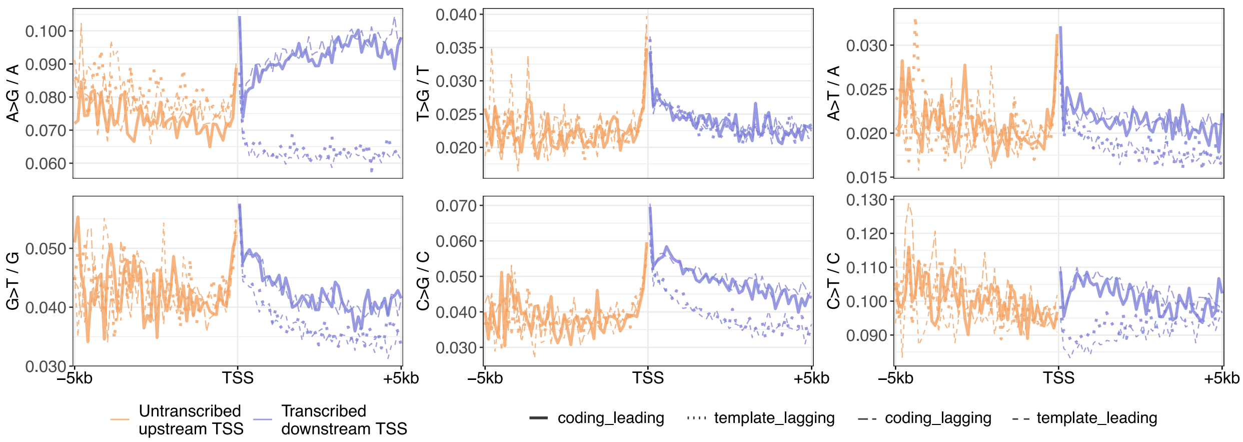


**Extended Data Figure 17:** non(CpG>TpG) ERV mononucleotide mutation density, stratified by transcription strand (coding or template) and replication strand (leading or lagging) in 100-bp windows around the TSS.
